# MLQ is responsible for stabilisation of subunit a in the holoenzyme of mammalian ATP synthase

**DOI:** 10.1101/2020.02.03.931709

**Authors:** K Tauchmannová, DH Ho, H Nůsková, A Pecinová, L Alán, M Rodinová, E Koňaříková, M Vrbacký, G Puertas, V Kaplanová, J Houštěk, P Pecina, T Mráček

## Abstract

The biogenesis of mammalian ATP synthase is complex process believed to proceed via several modules. It starts with the formation of F1 catalytic part, which is in the later steps connected with the membranous subcomplex. The final phase is represented by incorporation of the two mtDNA-encoded subunits Fo-a and A6L. However, little is known about the position of two newly described Fo accessory subunits DAPIT (also termed Usmg5) and MLQ (also known as c14orf2) in the assembly scheme and about their role in regulation of ATP synthase biogenesis. To resolve this, we have utilised several model systems, namely rho0 cells lacking mtDNA and thus both subunits Fo-a and A6L, cells harbouring 9205delTA microdeletion, which results in the absence of the subunit Fo-a, HEK293 cells with knockdown of DAPIT protein and HEK293 cells with knockout of MLQ protein and followed the assembly state of ATP synthase among them.

Contrary to previously reported data, we observed normal levels of assembled ATP synthase in DAPIT knockdown and MLQ knockout cells. Our results indicate that lack of DAPIT protein leads to the assembly of more labile, but complete and functional holoenzyme. Absence of either Fo-a alone or Fo-a and A6L results into the normal levels of structurally altered, labile, and ~60 kDa smaller vestigial enzyme complex, which also lacks DAPIT and MLQ. This complex retains the ATP hydrolytic activity but is unable to synthesize ATP. Cells with the MLQ knockout presented with the phenotype similar to the lack of Fo-a: normal content of smaller and labile complex. In the absence of MLQ, vestigial ATP synthase did not contain also subunits Fo-a and A6L. This complex also retained ATP hydrolytic activity, while its phosphorylating capacity was affected. In all the cell lines tested, the individual subunits seemed to be associated only with assembled ATP synthase complex, indicating that once subunits dissociate from the complex, they are degraded in the cell. This hypothesis is supported by the fact, that in the cells lacking subunit MLQ the biosynthesis of both mtDNA-encoded subunits Fo-a and A6L is normal, but they are degraded at faster pace than the rest of the complex.

Based on our data, we conclude that MLQ and Fo-a closely associate and their incorporation into the enzyme complex depends on each another. On the contrary, DAPIT protein seems to be incorporated at the very last step and its presence stabilises the holoenzyme.

## Introduction

Mitochondrial ATP synthase is central to cellular energy production, since it is responsible for the synthesis of up to 90% of ATP under normal physiological conditions. In addition to its primary function, ATP synthase can apparently play role in regulation of numerous other processes in mitochondria. Thus, ATP synthase oligomers are sequestered to the tips of mitochondrial cristae, where the constant angle between individual monomers contributes to the bending of the membrane and thus to establish cristae organisation (Strauss, Hofhaus et al. 2008, Davies, Strauss et al. 2011). Recently, ATP synthase was also demonstrated to be important component in the formation of mitochondrial permeability transition pore (mPTP), a large conductance channel in the inner mitochondrial membrane. Prolonged opening of mPTP collapses mitochondrial membrane potential (ΔΨ_m_) and ultimately leads to cell death. Finally, under certain conditions, ATP synthase can also reverse its function and sustain ΔΨ_m_ under conditions of respiratory chain blockade. This multifaceted nature of ATP synthase means that it is involved in regulation of vast number of cellular processes and thus faces multiple levels of regulation.

Principle of enzyme catalysis through rotary mechanism, together with solving of the structure of F1 part have been elucidated decades ago. However, detailed structure of the membrane intrinsic Fo part remained elusive until recently, when advances in cryoEM led to numerous high resolution structures from variety of organisms. While parts of the enzyme involved in the catalytic process, i.e. F_1_ domain, c-ring and proton half channels in the subunit A6L show remarkable degree of conservation, much greater level of diversity can be observed in the rest of F_o_ domain and especially for the peripheral stalk. Additional functions of ATP synthase, i.e. role in cristae formation or in mPTP formation are generally confined to these more divergent regions and may display greater variability across eukaryotic tree of life.

Since mitochondrial F_1_F_o_ ATP synthase is multisubunit enzyme, it is not surprising that its biogenesis requires ordered interaction between individual subunits in order to build-up the holoenzyme. For historical reasons, the current concept of the ATP synthase assembly is mostly based on yeast model, *S. cerevisiae* (Ackerman and Tzagoloff 2005, Rak, Zeng et al. 2009, Rak, Gokova et al. 2011), since only the recent advancements in genome editing opened the possibilities to study the mammalian complex in similar level of detail. The formation of ATP synthase from individual subunits was regarded as a stepwise process. It was shown to proceed via assembly of several modules, starting with an independent formation of F_1_ and oligomer of c-subunits. After the F_1_ attachment to the membrane-embedded c-ring, the subunits of peripheral arm (b, d, F_6_ and OSCP) and of membranous subcomplex (e, f, g, DAPIT and MLQ) are added. This model predicted that the two mtDNA-encoded subunits (a and A6L) are only added as the very last step of enzyme biogenesis (Wittig, Meyer et al. 2010). While this model still stands true in principle, recent studies on the mammalian enzyme challenged some of the assumptions. By systematically knocking out subunits e, f, g, DAPIT, MLQ (He, Ford et al. 2018) and c (He, Ford et al. 2017) Walker and coworkers demonstrated some level of variability regarding incorporation of individual modules. Thus, vestigial (and catalytically non-functional) complex can assemble without the c-ring, although the formation of F1-c module probably represents standard assembly intermediate. In these cells, it was also shown that the lack of MLQ protein decreases the amount of subunit F_o_-a and thus leads to defective assembly/stability of ATP synthase complex, that is accompanied with reduced respiration activity of mitochondria, while the absence of DAPIT protein have effect to the oligomers formation without any significant changes in respiration (He, Ford et al. 2018). Similarly, in fibroblasts of patients with mutation in Usmg5 gene, the major observed phenotype was reduced content of ATP synthase dimers, yet preservation of monomers (Barca, Ganetzky et al. 2018).

Conversely, recent advances in CryoEM imaging brought numerous new structures of ATP synthase and in particular its F_o_ region, which remained unresolved by classic crystallization techniques (Guo, Bueler et al. 2017, Muhleip, McComas et al. 2019). While good structures are still not available for F_o_ region of mammalian ATP synthase, the example of S. Cerevisiae shows that major dimerization surface is represented by subunit a (Guo, Bueler et al. 2017). The suspected S. cerevisiae ortholog of DAPIT – subunit k – would then be positioned laterally to the subunit a and have only small structural contribution to dimer interface. Given this information, it is surprising to observe major DAPIT KO phenotype to by lack of ATP synthase dimers (Barca, Ganetzky et al. 2018, He, Ford et al. 2018) and calls for detailed characterisation of dependencies between three last subunits incorporated during the enzyme assembly, namely subunit a, MLQ and DAPIT.

In the present work we used Clear-Native PAGE in combination with in-gel activity staining or WB antibody detection and revealed that ATP synthase lacking subunits Fo-a, A6L and MLQ is assembled in identical pattern with lower Mw (cV*). Absence of DAPIT did not influence incorporation of any other ATP synthase subunit. Rather surprising finding originated from pulse-chase studies, where we observed incorporation of subunit a into MLQ KO ATP synthase, thus rather than solely incorporation, the role of MLQ in in stabilisation of subunit a in the holoenzyme, since subunit a was quickly degraded, which resulted in observed steady-state pool of a-less ATP synthase in MLQ KO cells.

## Material and Methods

### Cell culture

Cells were cultivated under standard conditions (37 °C, 5% CO_2_ atmosphere) in the high-glucose DMEM medium (Life Technologies, 31966-021) supplemented with 10% fetal bovine serum (Life Technologies, 10270-106), 2 mM HEPES, and antibiotics (100 U/mL penicillin, 100 μg/mL streptomycin, Life Technologies, 15140-122). Cells were grown to ~90 % confluence, harvested using 0.05 % (w/v) trypsin and 0.02 % (w/v) EDTA, and washed twice in PBS before use.

### Cybrids and rho0

Transmitochondrial cybrids from patient harbouring 9205delTA mutation and from controls were prepared in the laboratory previously (Hejzlarova, Kaplanova et al. 2015). Heteroplasmy level was checked on a regular basis and was found to be >99%. Rho0 were osteosarcoma 143B TK- cells

### Models of *MLQ* knockout, DAPIT knockdown, AFG3L2 knockdown

*MLQ* knockout was generated in HEK293 cells by CRISPR-Cas9 variant employing the Cas9-D10A nickase guided by a pair of gRNAs complementary to both DNA strands of the targeted locus, essentially as described (Ran, Hsu et al. 2013). Briefly, HEK293 were co-transfected with a pair of pU6 vectors expressing gRNAs targeted to exon 2 of MLQ-coding gene *c14orf2* (Sigma-Aldrich custom product ID HSL0000631277 and HSR0000631282), along with pCMV vector expressing Cas9-D10A-GFP fusion protein (CAS9D10AGFPP, Sigma-Aldrich). Single cell clones were screened for a complete absence of MLQ protein by SDS-PAGE/WB and the deletion at the level of DNA was further pinpointed by Sanger sequencing of the region flanking the CRISPR-Cas9 cleavage site.

DAPIT knockdowns were generated in HEK293 cells using plasmids carrying pre-designed shRNAs (MISSION shRNA Plasmid DNA, Sigma-Aldrich) complementary to *ATP5MD* transcript. As a transfection reagent, Metafectene Pro (Biontex) was used in the ratio 2:1 to μg of plasmid DNA. Individual clones were isolated and kept under antibiotic selection (puromycin 1 μg/mL).

Knock-down of AFG3L2 in either control or MLQ KO HEK293 cells was performed by transient transfection with siRNA targeting *AFG3L2* transcript. Cells were transfected with specific siRNAs (Silencer Select s21518, Ambion) or non-silencing Negative Control siRNA, using Lipofectamine RNAiMAX (Life Technologies). Cells were transfected twice, at day 1 and day two post seeding, harvested 48h after the second transfection and frozen as dry pellets at −80°C for follow-up analyses.

### Electrophoresis and Western blot analysis

Samples for SDS-PAGE were denatured for 10 min at 60 °C in a sample lysis buffer containing 50 mM Tris-HCl pH 7.0, 4% (w/v) SDS, 10% (v/v) glycerol, and 0.1 M 1,4-Dithiothreitol and separated on a 10% or 12% polyacrylamide minigels (Bio-Rad MiniProtean III) using the Tricine buffer system (Schagger 2006) or using Any kD™ Mini-PROTEAN^®^ TGX precast gels (Bio-Rad #4569033) where indicated.

For native electrophoresis, tissue homogenates were centrifuged 10 min at 20000 *g* and cells 10 min at 600 *g*. The pellets were resuspended in buffer containing 50 mM NaCl, 2 mM 6-aminohexanoic acid, 50 mM imidazole, 1 mM EDTA, pH 7, solubilized with digitonin (2 g/g protein) for 20 min on ice and centrifuged for 20 min at 30 000 *g* to remove cell debris. Supernatants were differently treated for Blue-Native or Clear-Native Electrophoresis 1 (BNE, CNE) (Wittig, Carrozzo et al. 2007). Supernatants containing 10% glycerol and Coomassie Brilliant Blue G-250 (dye/detergent ratio 1:8) were analysed by BNE using 4-8% polyacrylamide gradient mini gels and the imidazole buffer system (Wittig, Braun et al. 2006). In case of CNE (Wittig and Schagger 2009) 10% glycerol and 0.01% Ponceau S were added to the supernatants. ATPase hydrolytic activity assay (Wittig, Carrozzo et al. 2007) was used to visualize active ATP synthase complexes in native gels.

For two-dimensional analysis, strips after 1^st^ dimension (BNE) were excised and proteins were denatured by 1h incubation in 1% SDS and 1% 2-mercaptoethanol. Gel slices were then subjected to SDS-PAGE using 10-16% gradient mini gels.

Western blot (WB) immunodetection was performed as previously (Hejzlarova, Tesarova et al. 2011). Membranes were blocked in 5% non-fat milk or 3% BSA, dissolved in TBS (Tris-buffered saline containing 150 mM NaCl, 10 mM Tris-HCl, pH 7.5) for 1 h prior to incubations with primary (2 h at room temperature or overnight at 4 °C) antibodies to subunits of ATP synthase (F_1_-α (Moradi-Ameli and Godinot 1983) lot 20D6, F_1_-β (Abcam ab14730), F_1_-γ (GTX 114275), F_1_-*δ* (GTX101503), F_o_-a (Dubot, Godinot et al. 2004), F_o_-b (Abcam 117991), F_o_-c (Abcam 181243), F_o_-d (GTX 87685), F_o_-g (GTX 111014), A6L (Biorbyt orb215488), OSCP (Santa Cruz SC-98707), DAPIT (Proteintech 17716-1-AP), MLQ (Proteintech 14704-1-AP), Complex II (SDHA, Abcam 14715), AFG3L2 (Proteintech 14631-1-AP), OPA1 (BD Bio 612606), and actin (Calbiochem CPO1-1EA). For a quantitative detection, the corresponding infra-red fluorescent secondary antibodies (Alexa Fluor 680, Life Technologies; IRDye 800, LI-COR Biosciences) were used. The fluorescence was detected using Odyssey infra-red imaging system (LI-COR Biosciences) or ChemiDoc Imaging System (Bio-Rad) and the signal was quantified using Aida 3.21 Image Analyzer software (Raytest).

### Metabolic pulse-chase labeling of mtDNA encoded proteins

Proteins encoded by mtDNA were labeled using ^35^S-Protein Labeling Mix (Met+Cys; Perkin Elmer NEG072) by procedure described in (McKenzie, Lazarou et al. 2009). Briefly, cells were incubated for 16h with chloramphenicol (40 μg/ml), washed twice in PBS, and after 15 min incubation in DMEM medium without methionine and cysteine (DMEM-Met-Cys) and 15 min incubation in DMEM-Met-Cys with cycloheximide (CHX; 0.1 mg/ml), ^35^S-Protein Labeling Mix (350 μCi/150 mm dish) was added. Cells were incubated for 3h, then 250 μM cold Met and Cys was added and after 15 min cells were washed with PBS + 250 μM cold Met and Cys and finally with PBS. Cells were grown in standard DMEM medium supplemented with 5% (v/v) fetal bovine serum and harvested at different times (24,48,72,96 hrs). Pellets of labeled cells were mixed with the same w/w of unlabeled cells, cell membranes were isolated, solubilized with digitonin and analyzed by 2D BNE/SDS PAGE. Gels were stained in a Coomassie R 250 dye, dried and radioactivity was detected using Pharos FX™ Plus Molecular Imager (Bio-Rad).

### Immunoprecipitation

Cells were harvested, solubilized with digitonin 2 g/g protein in PBS containing PIC (1:500, Sigma P8340) for 10 min on ice and centrifuged at 30 000 *g* for 15 min. Supernatant was incubated with anti-ATP synthase immunocapture kit (Abcam ab109715) on a nutator for 5 h at 4 °C. The resin was washed 2x with PBS + PIC + 0.05% digitonin and 2x with PBS + PIC. Bound proteins were eluted by SDS sample buffer and analyzed by SDS-PAGE and WB.

### LFQ protein mass spectrometry analysis

100 μg of cell pellets (HEK293, MLQ KO, DAPIT KD) were solubilized using sodium deoxycholate, reduced with TCEP [tris(2-carboxyethyl)phosphine], alkylated with MMTS (S-methyl methanethiosulfonate), digested sequentially with Lys-C and trypsin, and extracted with ethylacetate saturated with water as described (Masuda, Tomita et al. 2008). Samples were desalted on Empore C18 columns, dried in Speedvac and dissolved in 0.2% TFA. 1 μg of peptide digests were separated on 50 cm C18 column using 2.5 h elution gradient and analyzed on Orbitrap Fusion Tribrid (Thermo Scientific, USA) mass spectrometer. Resulting raw files were processed in MaxQuant (v. 1.5.3.28) (Cox and Mann 2008) with label-free quantification (LFQ) algorithm MaxLFQ (Cox, Hein et al. 2014). Downstream analysis and visualization were performed in Perseus (v. 1.5.3.1).

### Electron microscopy

For electron microscopy analysis, the cells were fixed according to (PMID: 13398447) with slight modifications. Briefly, the cells were incubated in PBS containing 2% potassium permanganate for 15 min, washed with PBS, and dehydrated with an ethanol series. The cells were subsequently embedded in Durcupan Epon (Electron Microscopy Sciences, Hatfield, PA, USA), sectioned by Ultracut microtome (Reichert, Depew, NY, USA) to thicknesses ranging from 600 to 900 Å, and finally stained with lead citrate and uranyl acetate. A JEOL JEM-1200 EX transmission electron microscope (JEOL, Tokyo, Japan) was used for image acquisition.

### Oxygen consumption and extracellular acidification

The Seahorse XF^e^24 Analyzer (Agilent) was used to measure oxygen consumption and extracellular acidification of intact cells that were seeded on poly-L-lysine (Sigma-Aldrich) coated plates on a day before the measurement (30,000–40,000 cells per well). The measurements were carried out in the XF base medium supplemented with 10 mM glucose, 1 mM pyruvate, 2 mM L-glutamine, and 0.2 % bovine serum albumin. Final concentrations of applied inhibitors were as follows: 1 μM oligomycin, 2 μM FCCP (carbonyl cyanide-p-trifluoromethoxyphenylhydrazone), 1 μM rotenone, 0.5 μM antimycin A, and 100 mM 2-deoxyglucose. Both oxygen consumption and extracellular acidification were normalized to the cell count estimated after nuclear staining with Hoechst on BioTek Cytation 3 imaging plate-reader (Agilent, USA). Image analysis and cell count was performed in Gen5 Image+ software.

## Results

### Characterisation of the cells lacking MLQ or DAPIT proteins

To study the functional role of MLQ and DAPIT proteins in more detail, we established HEK293 cell lines with knockout of MLQ (MLQ KO, using CRISPR/CAS9), and with knockdown of DAPIT protein (DAPIT KD, using shRNA). To characterise the cells from the bioenergetics point of view, we analysed basic mitochondrial functions of the cells (Figure 1). Using Seahorse instrument we measured OCR and ECAR activities of the cells in a medium with TCA cycle substrates glutamine and pyruvate.

**Figure 1:**
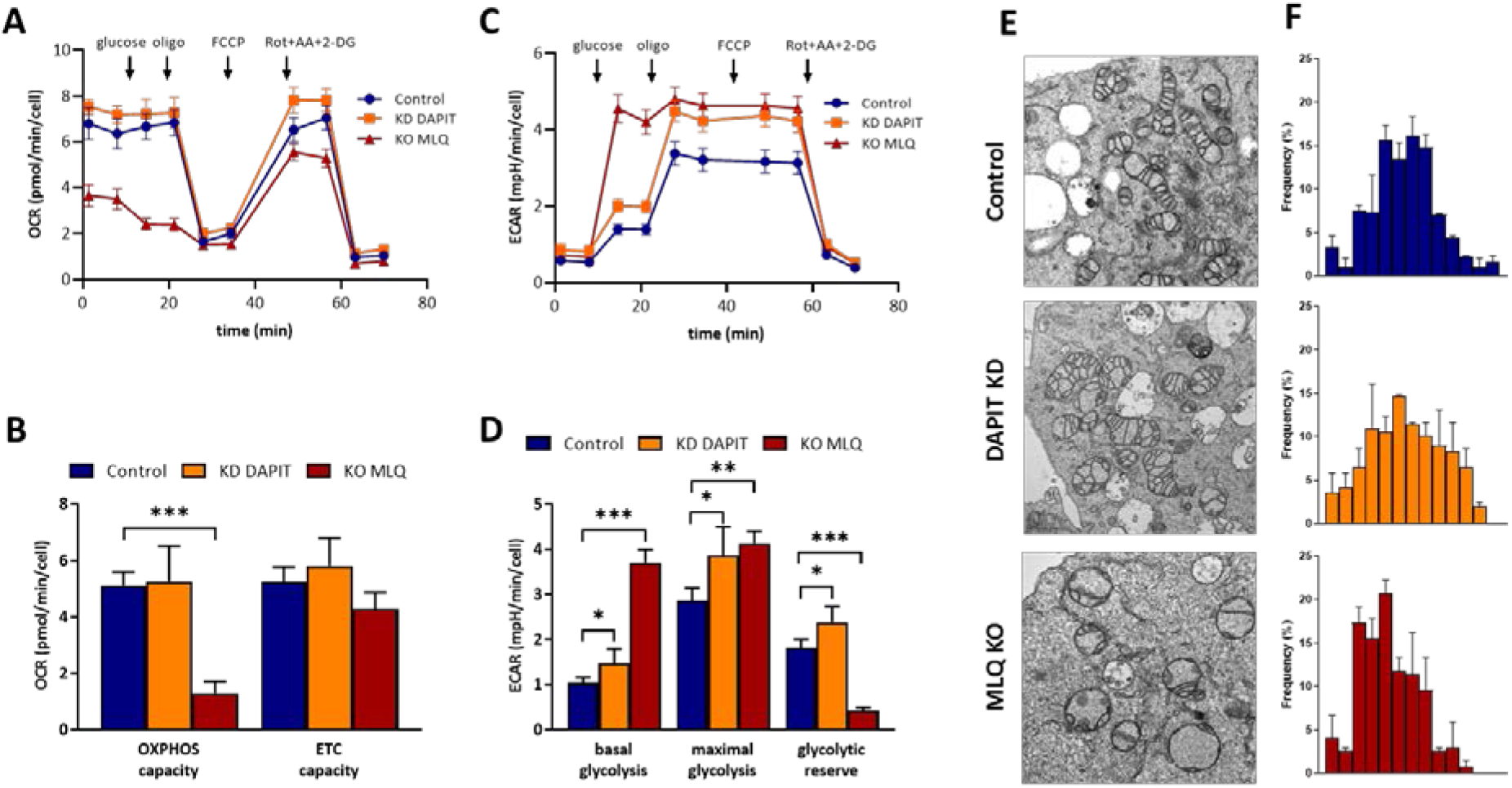
Respiratory function and structure of mitochondria in the cells lacking MLQ or DAPIT protein. Seahorse analysis of mitochondrial respiration (**A, B**) and glycolytic rate (**C, D**) in MLQ KO and DAPIT KD cells was performed in a medium with TCA cycle substrates glutamine and pyruvate. OXPHOS and ETC capacities were calculated from OCR activities, basal and maximal glycolysis and glycolytic reserve were evaluated from ECAR activities; oligo – oligomycin, Rot – rotenone, AA – antimycin A, 2-DG – 2-deoxyglucose, values are mean ± SD, n=3-4, statistics - * p<0.05, ** p<0.01, *** p<0.001. **(E)** Electron microscopy analysis was used to calculate frequency distribution of relative cristae length per surface area of mitochondria (**F**) in of control HEK (blue), MLQ KO (red) and DAPIT KD (orange) cells.

We found that in cells lacking MLQ the basal respiration corresponding to the effectivity of the whole OXPHOS system was very low compared to control and also failed to increase O_2_ consumption after OXPHOS stimulation by uncoupler (Fig. 1 A,B). Conversely, addition of glucose during glycolysis stress protocol revealed that MLQ KO cells were more likely to shift towards glycolysis than control cell line, as seen by more pronounced increase in the extracellular acidification rate (ECAR, dependent directly on the glycolytic activity). Interestingly, increased glycolytic rate manifested in MLQ KO cells also as decrease in the oxygen consumption rate (OCR), i.e. Crabtree effect (Fig. 1 C,D). This Crabtree effect was much less prominent in control cells which preferred to obtain energy from respiration instead glycolysis. For more, MLQ KO cells are more glycolytic at the basal level. In contrast, the respiring efficiency of cells lacking DAPIT protein was much less affected (Fig. 1 A,B) and also their basal glycolysis and glycolytic reserve differed less from the controls (Fig. 1 C,D).

Transmission electron microscopy of mitochondrial ultrastructure in MLQ KO and DAPIT KD revealed that the energetic dysfunction was also associated with ultrastructural changes in MLQ KO, which displayed disturbed mitochondrial ultrastructure with swollen mitochondria and less cristae pre length (Fig. 1E). These changes were also assessed quantitatively, as compared to controls, MLQ KO mitochondria (red bars) clustered towards the left side of the graph – lower relative length of cristae per surface area of mitochondria. No similar changes could be found in DAPIT KD cells (orange bars).

### Content of ATP synthase subunits

In the next step we analysed the changes in the cellular content of ATP synthase subunits by WB immunodetection and LFQ MS analysis, to assess the structural consequences of MLQ and DAPIT absence, in particular with respect of the mtDNA-encoded subunits F_o_-a and A6L (ATP6 and ATP8). For comparison, we also included mammalian cellular models, where the mtDNA-encoded ATP synthase subunits are affected. Specifically, these involved 143B TK^+^ (ρ^+^) and 143B TK^−^ (ρ^0^) osteosarcoma cells with or without mtDNA, respectively, and cybrid cell line with homoplasmic mtDNA m.9205delTA mutation (ΔTA) in *MT-ATP6* gene (Hejzlarova, Kaplanova et al. 2015) which prevent expression of ATP6/8 genes as models of F_o_-a and A6L deficiency. Isolated mitochondria from the cell cultures were analysed by SDS-PAGE and WB, and quantitative detection of these subunits as well as other F_o_ and F_1_ subunits was performed (Figure 2). The efficacy of the primary depletion of DAPIT and MLQ was almost 100 % and equally effective was depletion of F_o_-a and A6L in ρ^0^ and ΔTA cells. In accordance with published data (He, Ford et al. 2018) we have found that absence of MLQ results in the decrease of F_o_-a, A6L and DAPIT while absence of DAPIT has no effect on other subunits. On the other hand, in ρ^0^ cells we observed marked decrease of MLQ and even more pronounce decreased of DAPIT, and similar changes were present in ΔTA cells. Contrary to originally reported data (Ohsakaya, Fujikawa et al. 2011, Fujikawa, Ohsakaya et al. 2014) and in accordance with the recent study (Walker et al 2018) neither MLQ knockout, nor DAPIT knockdown displayed profound ATP synthase deficiency, since the remaining ATP synthase subunits are present in normal amounts (Fig. 2).

**Figure 2:**
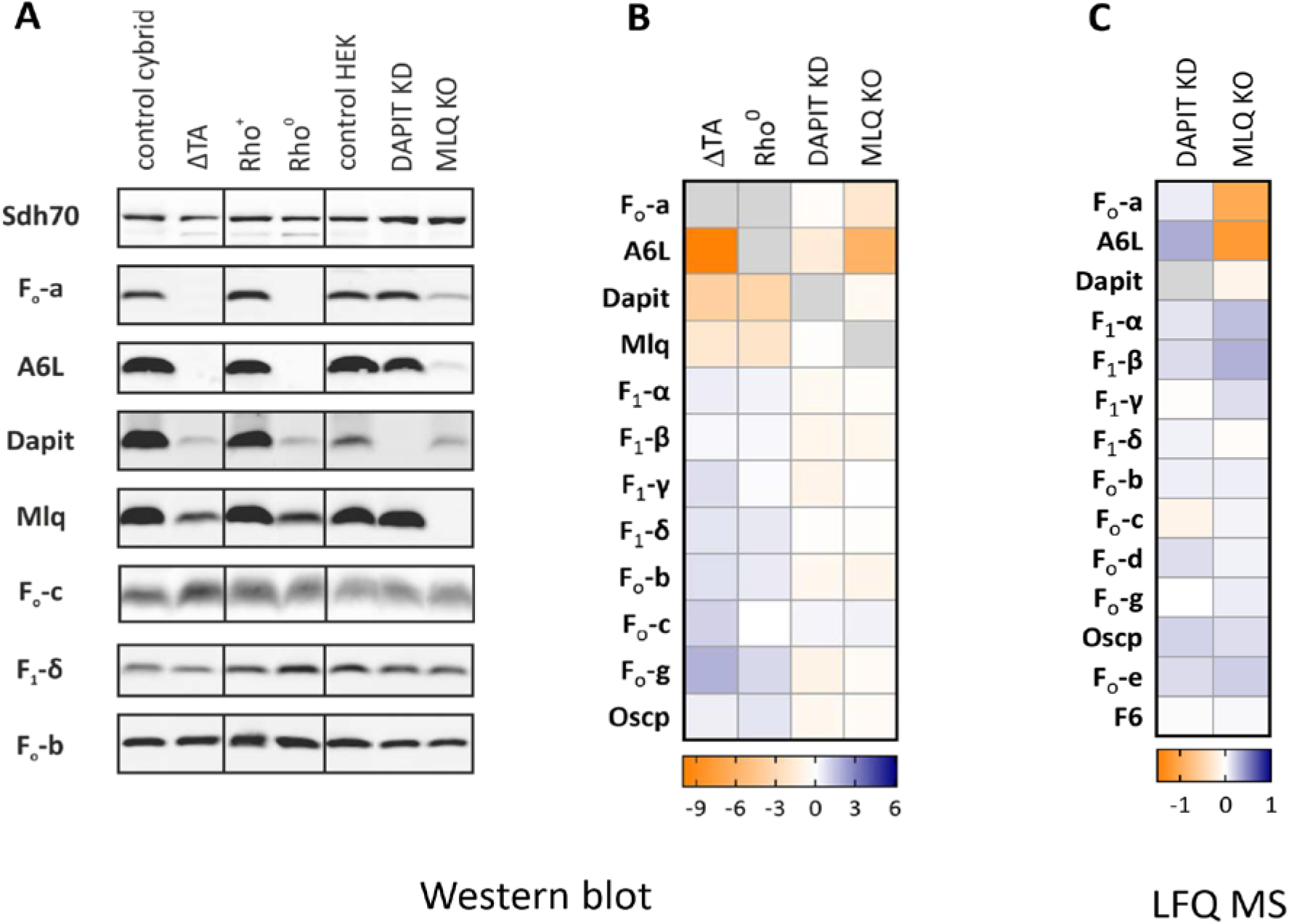
The content of ATP synthase subunits in MLQ KO and DAPIT KD cells. **(A)** SDS-PAGE and WB image of immunodetected subunits in isolated mitochondria of MLQ KO, DAPIT KD, Rho^o^ and ΔTA cells. **(B)** WB signal of ATP synthase subunits was normalised to Sdh70 subunit of respiratory chain complex II, quantification displays mean values of the subunit content fold-change relative to respective control. **(C)** LFQ MS/MS quantification of ATP synthase subunits in MLQ KO and DAPIT KD cells.

### Native forms of ATP synthase

Further, we were interested to assess by native electrophoresis how the lack of the respective subunits affects the assembly of ATP synthase and the association of the other subunits into the enzyme complex. To maintain the native situation in the mitochondrial membrane and to avoid breakup of ATP synthase during electrophoretic analysis, we used very mild conditions for separation of ATP synthase complex. Mitochondria from respective cell lines were solubilised by digitonin 2 g/g protein, and separated on clear native PAGE (CNE) without Coomassie Blue dye. ATP synthase was then visualised in gel by ATP hydrolytic activity staining or by WB detection with specific antibodies (Figure 3). The overall amount of ATP synthase was unchanged in all the cell lines tested and the enzymes retained their hydrolytic activity. In-gel ATPase activity showed the presence of dimers and oligomers in MLQ KO and DAPIT KD, they were also detected in ΔTA and Rho^0^ cells but their size was smaller (**cV**_**D**_**\***, **cV**_**o**_**\***). Furthermore, MLQ KO, Rho^0^ and ΔTA cells displayed shift from oligomers to dimers and monomers. All ATP synthase supercomplexes were clearly visible by WB with F_1_-α antibody, while their detection with DAPIT, MLQ, F_o_-a and A6L antibodies mirrored the subunit expression as shown above in Figure 2. Thus, ATP synthase complexes in MLQ KO cells lacked MLQ, contained only traces of F_o_-a and A6L, and also displayed decreased content of DAPIT. To the contrary, ATP synthase of DAPIT KD only lacked DAPIT protein. As expected, ATP synthase complexes in Rho^0^ and ΔTA lacked Fo-a and A6L, but also MLQ and DAPIT.

**Figure 3:**
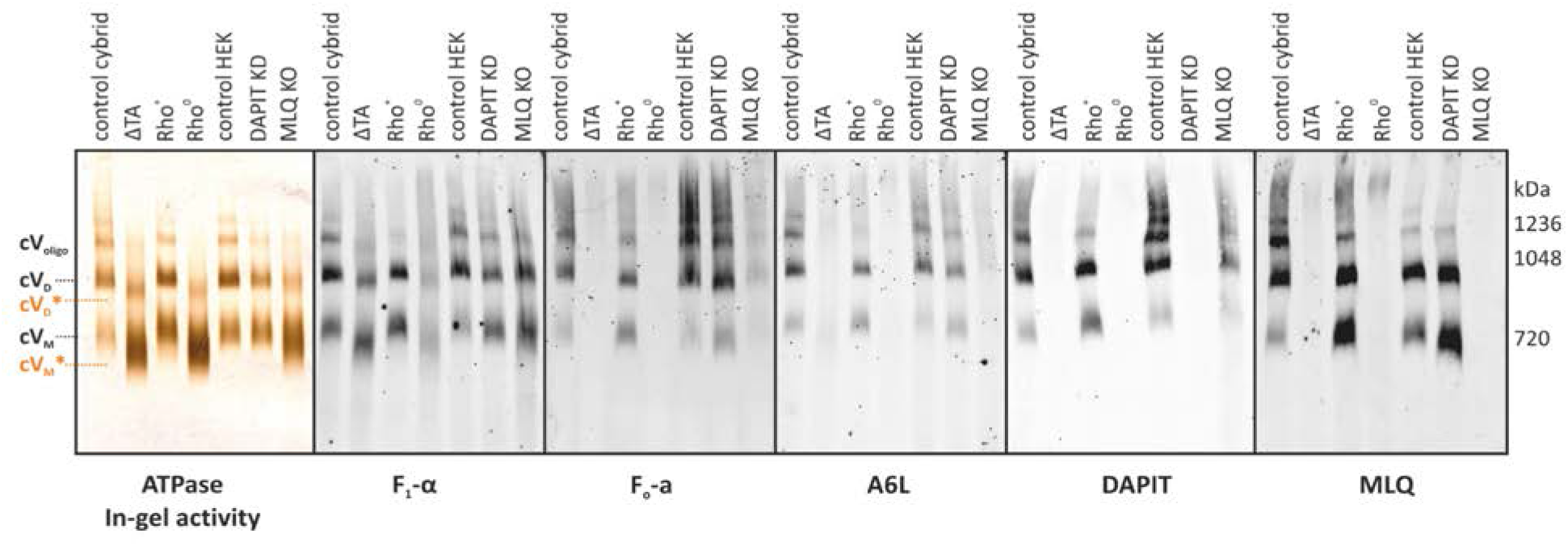
Native forms of mitochondrial ATP synthase. Mitochondria isolated from cell lines were solubilised by digitonin 2 g/g protein, separated on clear-native PAGE (CNE) and either stained in gel for ATP hydrolytic activity or used for WB detection of F1-α, Fo-a, A6L, DAPIT and MLQ subunits. Individual forms of ATP synthase are indicated: cV_M_ – monomer, cV_D_ – dimer, cV_o_ – oligomer, cV_M_* and cV_D_* denote smaller form of the respective complexes.

In ΔTA and Rho^0^ cells smaller dimeric (cV_D_^*^) and monomeric (cV_M_^*^) forms of ATP synthase were detected, which is in accordance with previously published data (Hejzlarova, Kaplanova et al. 2015). The difference in size, previously estimated as 50-60 kDa, is in good agreement with the missing F_o_-a+A6L+DAPIT+MLQ (25+8+7+7 kDa). Smaller forms of ATP synthase can be seen also in MLQ KO; here the signal of monomer spreads down, possibly due to the co-existence of fully assembled complex containing the remaining amount of subunits F_o_-a, A6L and DAPIT and the vestigial complex without these subunits.

### Stability of ATP synthase complex

The above CNE profiles of ATP synthase indicated possible alterations in supercomplex stability. It is well known, that less stable ATP synthase complexes can be disintegrated during separation at harsher/more stringent conditions of electrophoresis such as upon exposure to Coomassie Blue Dye (Wittig, Meyer et al. 2010, Hejzlarova, Kaplanova et al. 2015, Kovalcikova, Vrbacky et al. 2019). Thus, BNE was further used to assess stability of ATP synthase complexes (Figure 4). As expected, the lack of subunits F_o_-a and A6L results in the decreased stability of ATP synthase complex, since almost no dimers (cV_D_) or monomers (cV_M_) are found in Rho^0^ and ΔTA cells, where the majority of ATP synthase complex is broken down to F_1_ containing subcomplexes (mainly F_1_-c).

**Figure 4:**
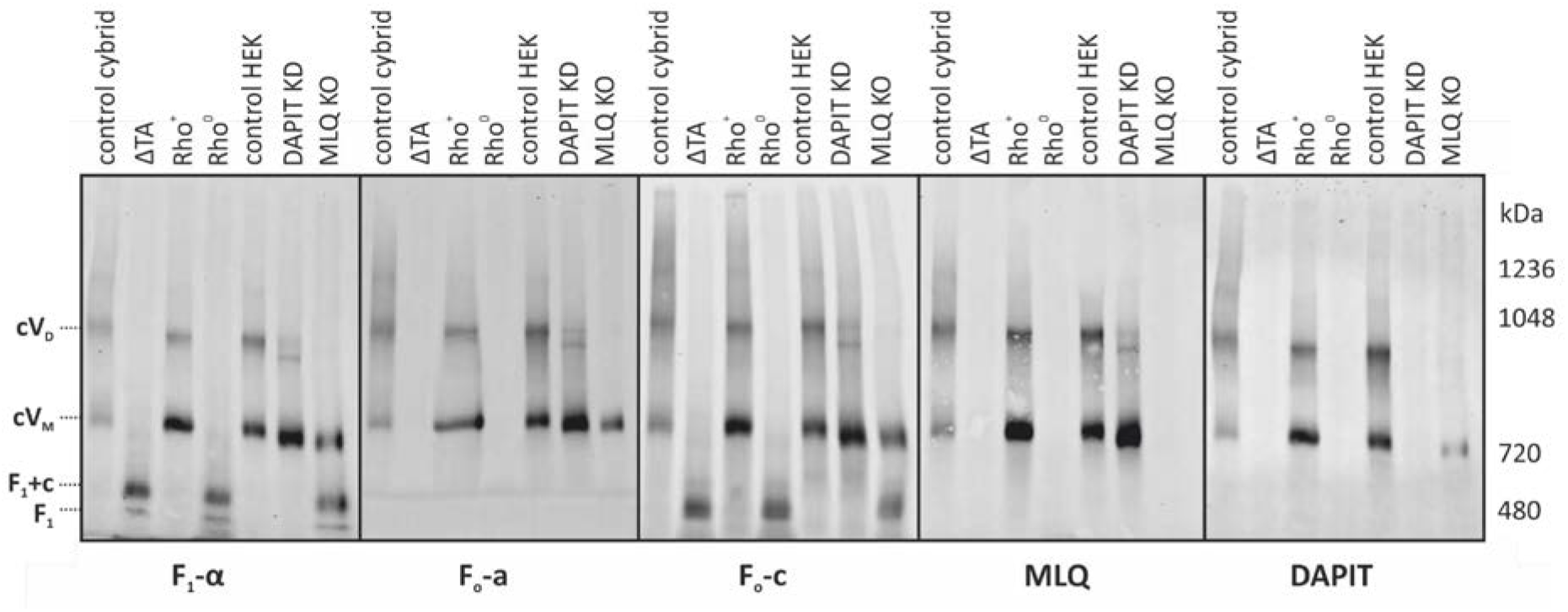
Absence of F_o_-a / MLQ yields labile holoenzyme, which dissociates into F1-c and F1 intermediates under the conditions of BNE electrophoresis. Mitochondria isolated from the respective cell lines were solubilised by digitonin 2 g/g and separated on blue-native PAGE (BNE). WB detection of F1-α, Fo-a, A6L, DAPIT and MLQ subunits. ATP synthase forms: cV_M_ – monomer, cV_D_ – dimer, F_1_+c − F_1_ and c subunits and F_1_ subcomplexes.

**Figure 5:**
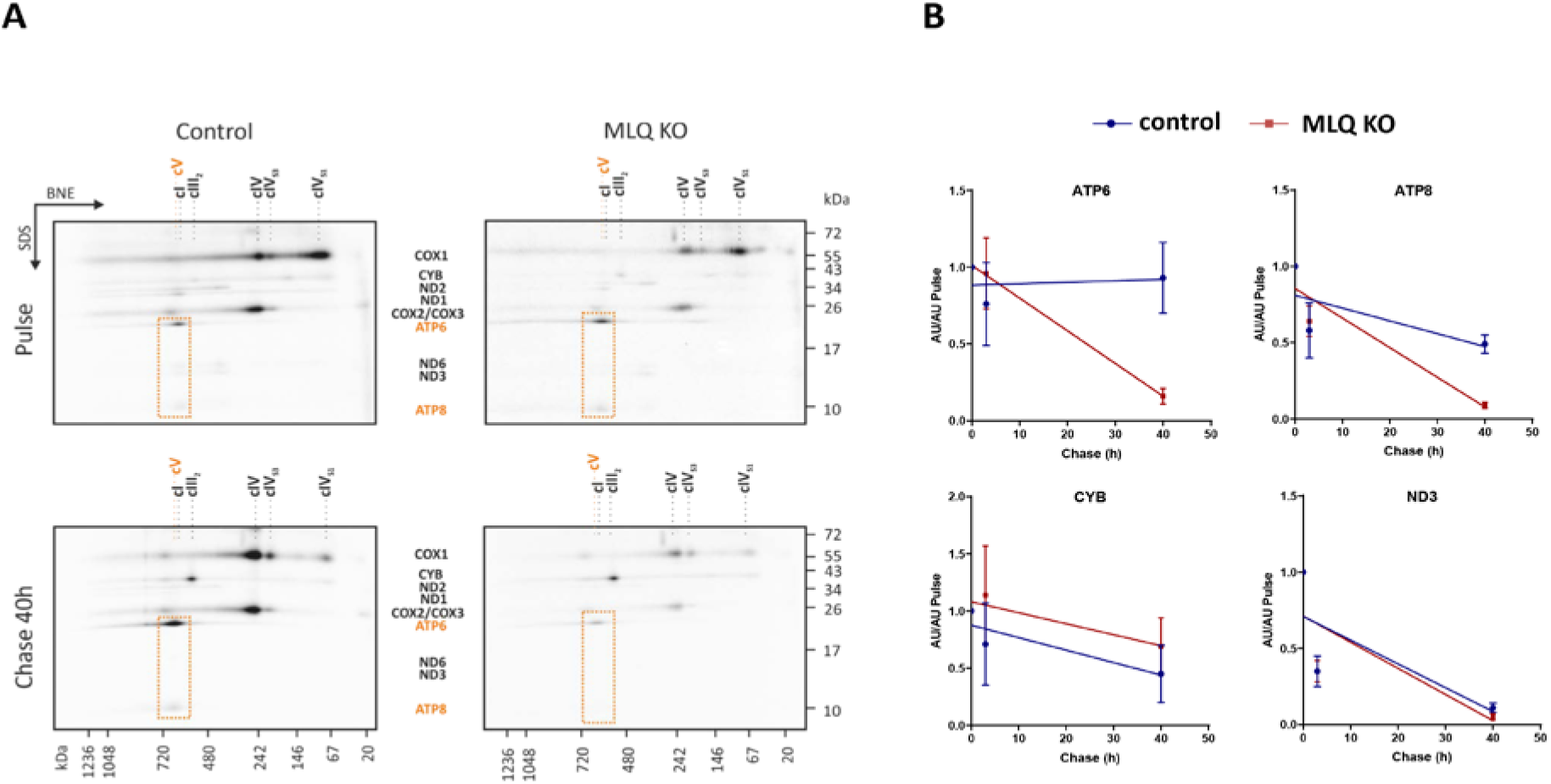
Rapid degradation of mtDNA encoded subunits in MLQ knockout. (**A**) mtDNA-encoded subunits were radioactively labelled with ^35^S-methionin/cysteine and subsequently chased for 40h with cold methionine. Cells were solubilised with digitonin and analysed using 2D BNE/SDS PAGE. (**B**) Quantification of radioactivity of selected subunits F_o_-a (ATP6), A6L (ATP8), CYT B and ND3 based on the separation in SDS-PAGE gel.

In MLQ KO, resting amount of subunit F_o_-a and A6L associated with ATP synthase maintained the stability of the complex lacking only MLQ, while the incomplete ATP synthase, lacking both MLQ and F_o_-a behaves in the same way as in Rho^0^ and ΔTA cells.

In DAPIT KD, smaller monomeric form of ATP synthase complex is present. This complex only lacks DAPIT protein, shows the same stability as ATP synthase in control cells and does not disintegrate to F_1_ subcomplex. The overall amount of normal sized dimeric form is significantly decreased, and a population of ATP synthase with lower mass than dimer but higher than monomer is present. Again, this complex seems to lack only DAPIT protein. In conclusion, decrease in the content of higher oligomers and increase in monomer (most likely arising from decomposed oligomers) suggests the decreased stability of ATP synthase oligomers in DAPIT KD cells.

### Turnover of mtDNA encoded subunits in MLQ KO

From previous experiments we can conclude, that absence of MLQ protein result in severe reduction of subunits F_o_-a and A6L levels, but the resting amount of the subunits is still associated with ATP synthase complex, suggesting that synthesis of mtDNA encoded subunits or its assembly into ATP synthase is affected. To resolve this, we performed radioactive metabolic labelling of mtDNA encoded proteins in MLQ KO cells and labelled proteins were analysed by SDS and 2D BNE/SDS-PAGE (Fig. 6, S3). For the pulse, cells were labelled for 3 hours, then it was grown in normal culture media for 3 hours (chase 3 h) or 40 hours (chase 40 h). Surprisingly, both F_o_-a (ATP6) and A6L (ATP8) subunits were synthesised and assembled into ATP synthase, since in the pulse sample the amount of labelled subunits associated with the complex was normal or even higher than in control cells (Fig. 6A). After short time in culture (chase 3 h), the amount of assembled subunits was still similar in both control and MLQ KO cells without any significant degradation (Fig. S3). However, after 40 hours in culture (Chase 40 h) the amount of labelled subunits assembled into ATP synthase was distinctly increased in the control cells while in MLQ KO we can see significant decrease in the amount of F_o_-a and A6L subunits associated with ATP synthase complex. In other words, F_o_-a and A6L are compelled to faster degradation in K/O MLQ cells. (Higher synthesis and) faster degradation of these subunits in MLQ KO is nicely demonstrated also in Fig. 6B, which shows the quantification of labelled mtDNA-encoded proteins from normal SDS-PAGE (Fig. S3), thus reflecting the overall amount of the newly synthesized subunits. This phenomenon was specific for ATP synthase subunits, since the profile of other respiratory chain subunits was similar in HEK and KO cells, for example of subunits ND3 and ND6 (Fig. 6).

**Figure 6:**
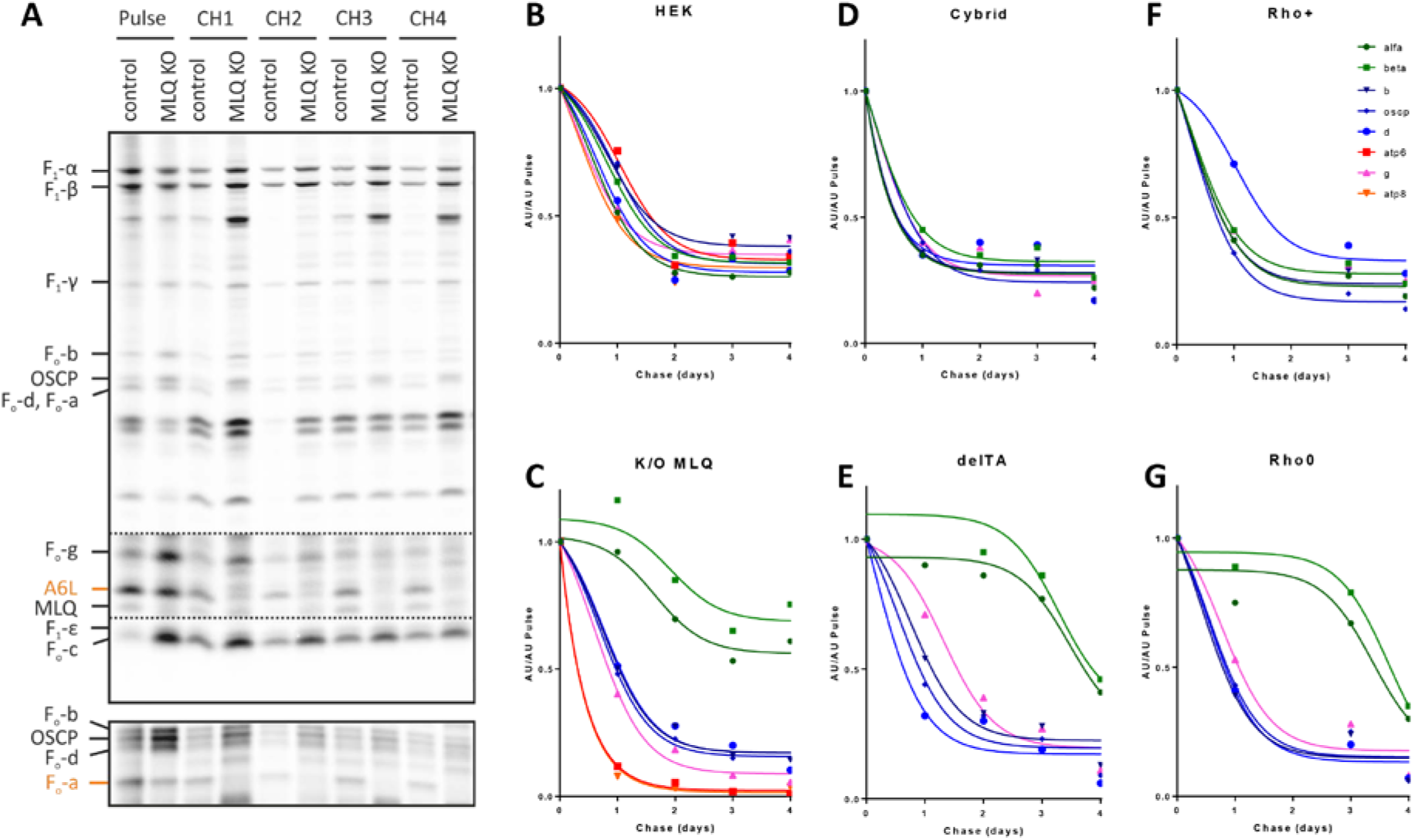
Modular kinetics of ATP synthase assembly into holoenzyme. **(A)** MLQ KO and control cells were radioactively labelled for 3h with ^35^S-methionin/cysteine and subsequently chased with cold methionine for 1, 2, and 3 days. Cells were harvested, solubilised with triton X-100 and ATP synthase was isolated by immune-affinity purification using ATP synthase immunocapture kit. IP-ATP synthase was than analysed by SDS-PAGE. Quantification of the radioactive signal of all ATP synthase subunits shows uniform kinetics of degradation in control HEK cells **(B)**, while **(C)** MLQ KO cells displayed faster turnover of subunits F_o_-a and A6L (red lines) but stabilization of the F_1_ subunits (α and β, green lines) compared to subunits b, d and OSCP of the external stalk (blue line). Similar stabilisation of F_1_ subunits show ΔTA cells (**E**) compared to control cybrids (**D**) as well as rho^0^ cells (**G**) compared to control rho^+^ cells (**F**).

To characterize the turnover of individual ATP synthase subunits in MLQ KO cells in more detail, we then analysed the metabolically labelled proteins in ATP synthase complexes isolated by immunoprecipitation. MLQ KO and control HEK cells were radioactively labelled for 3 hrs and subsequently chased for 1, 2, and 3 days. Cells were harvested, solubilised with Triton X-100 and ATP synthase was isolated by immune-affinity purification using ATP synthase immunocapture kit. Labelled subunits of IP-ATP synthase complexes were resolved by gradient SDS-PAGE (Fig. 7A). Turnover of ATP synthase subunits showed similar turnover of all subunits in control cells with a T1/2 of XX-XX days (Fig. 7B). In contrast, in MLQ KO their decay and turnover largely differed and manifested in 3 groups (Fig. 7c). The peripheral stalk subunits Fo-b, OSCP and Fo-d behaved similarly as subunits in control cells while the other subunits strongly differed. On one hand, the Fo-a subunit was much faster degraded, on the other hand the F_1_ subunits α, β and γ displayed much slower degradation and longer half-life. Absence of MLQ thus caused a rapid degradation of mtDNA encoded subunits initially attached to the ATP synthase complex thus leading to a steady state ATP synthase complex almost devoid of Fo-a and A6L. Interestingly, this led also to a stabilisation of F_1_ catalytic part. Overall, upon MLQ deficiency, ATP synthase showed a modular behaviour with respect of degradation of primarily assembled subunits.

## Discussion

This paper focuses on the interdependencies between four subunits of ATP synthase – ATP6, A6L, DAPIT and MLQ in their assembly into the enzyme. While initial studies of Yoshida group suggested severe reduction in the content of assembled ATP synthase for cells lacking DAPIT or MLQ, this view has been challenged by recent literature, which suggested role of DAPIT only in enzyme dimerization and requirement of MLQ for proper assembly of ATP6 into the holoenzyme.

Assembly of mammalian mitochondrial ATP synthase complex is still not fully understood. One part of the process that is the subject of recent discussion is the assembly of small subunits of the F_o_ part, mitochondrially encoded F_o_-a and A6L subunits, and MLQ and DAPIT proteins, that were described as ATP synthase subunits few years ago (Meyer, Wittig et al. 2007, Ohsakaya, Fujikawa et al. 2011, Fujikawa, Ohsakaya et al. 2014). Recently, Walker et al. showed (He, Ford et al. 2018), that in the cells lacking Dapit or MLQ proteins, the respiration is slightly or severely decreased, respectively, but enhanced oligomycin sensitivity, while in the studies of Yoshida group, cells lacking both Dapit or MLQ produce ATP in much lower levels than control cells. Also the effects to ATP synthase assembly was not consistent in these models.

Both MLQ and DAPIT are highly basic (pI~10) small (MW~8 kDa) proteins with one putative transmembrane domain. They co-purify with ATP synthase only if it is isolated in the presence of exogenous phospholipids (Chen, Runswick et al. 2007) or after solubilisation with mild detergents (Meyer, Wittig et al. 2007). There are some indications that both DAPIT (Ohsakaya, Fujikawa et al. 2011) and MLQ (Fujikawa, Ohsakaya et al. 2014) knockdowns in cell lines lead to decrease in the content of assembled ATP synthase, but other studies contradict this (Baughman, Nilsson et al. 2009). Another hint about potential function of MLQ and DAPIT may come from the observation, that ATP synthase isolated in coupled state displayed higher resistance to limited proteolysis (Runswick, Bason et al. 2013). This implicates change in enzyme conformation between coupled and uncoupled states. Of interest here is, that conditions of isolation, which yield coupled enzyme are the same as those retaining DAPIT and MLQ in enzyme structure, i.e. presence of phospholipids and especially cardiolipin.

As both MLQ and DAPIT easily dissociate from the complex, they are supposedly assembled at the later stages of ATP synthase biogenesis. We hypothesize that late assembly position imposes them to be involved in incorporation or stabilization of mitochondrial encoded subunits Fo-a and A6L that are proposed to be the last subunits assembled into mammalian ATP synthase. To resolve this, we followed the assembly state of ATP synthase in the several cellular model systems with deficient expression of implicated subunits. We further studied how MLQ absence affects the structure and function of the enzyme and whether the defect influenced the respiratory chain.

Similarly to respiratory chain also ATP synthase organises into higher order assemblies: it is now widely accepted that ATP synthase from mitochondria is present in the membrane in dimers (Wittig and Schagger 2009, Wittig, Meyer et al. 2010). Interaction between two monomers occurs via F_o_ and appears to involve subunits a, e, g, b and A6L. While this is the situation in yeast, in mammals dimerization may be a more complex process as IF1 interacting with F_1_ was also proposed to mediate it (Campanella, Casswell et al. 2008). However, even here the physical releasing of IF1 from the inner mitochondrial membrane did not alter the amount of dimers extracted by mild detergent (Tomasetig, Di Pancrazio et al. 2002). As the monomers interact with each other at 70–90° angle and the neighbouring dimers at 20° angles, the supramolecular ATP synthase ribbons can shape the inner membrane at the apical regions and thus support cristae formation. In addition, a recent numerical simulation indicates that such supramolecular assembly may favour effective ATP synthesis under proton-limited conditions. This means that assembly into supercomplexes may dictate efficiency of ATP production analogously to the optimisation of electron flux through OXPHOS supercomplexes (Lapuente-Brun, Moreno-Loshuertos et al. 2013).

We demonstrate that binding of MLQ to cV assembly intermediate precedes attachment of ATP6 and A6L, in its absence MLQ binding is unstable and it falls off. Structurally, MLQ KO ATP synthase can form dimers and even oligomers, but these supercomplexes are unstable. As a consequence, also the structure of mitochondria is affected, since it presents with shorter cristae and swollen mitochondria Functionally, MLQ KO ATP synthase cannot synthesize ATP and energy metabolism switches to glycolysis. Overall, MLQ deficiency leads to serious structural and functional impairment of ATP synthase and thus mitochondrial energy provision indicating that MLQ mutations may lead to mitochondrial disease

## Abbreviations

F_1_: catalytic part of ATP synthase
F_o_: membrane-embedded part of ATP synthase
ΔΨ_m_: membrane potential of mitochondrial membrane
ROS: reactive oxygen species
mtDNA: mitochondrial DNA

## Acknowledgements

This work was supported by the Grant Agency of the Czech Republic (16-01813S) and Ministry of Health of the Czech Republic (16-33018A). Institutional support was provided by RVO: 67985823.

---

**Table 1:** cell models used in the study.

| Cell type | Missing subunit | Designation |
| --- | --- | --- |
| Cybrid cells with wild type mtDNA | None | Control cybrid |
| Cybrid cells with homoplasmic mtDNA m.9205delTA mutation in *MT-ATP6* gene | Fo-a, A6L | ΔTA |
| Control osteosarcoma cell | None | Rho<sup>+</sup> |
| Osteosarcoma cells without mtDNA | Fo-a, A6L | Rho<sup>0</sup> |
| HEK293 cells | None | Control |
| HEK293 cells with shRNA knockdown of *ATP5MD* gene | DAPIT | DAPIT KD |
| Hek293 cells with CRISPR/Cas9 knockout of *MP68* gene | MLQ | MLQ KO |

**Figure S1:**
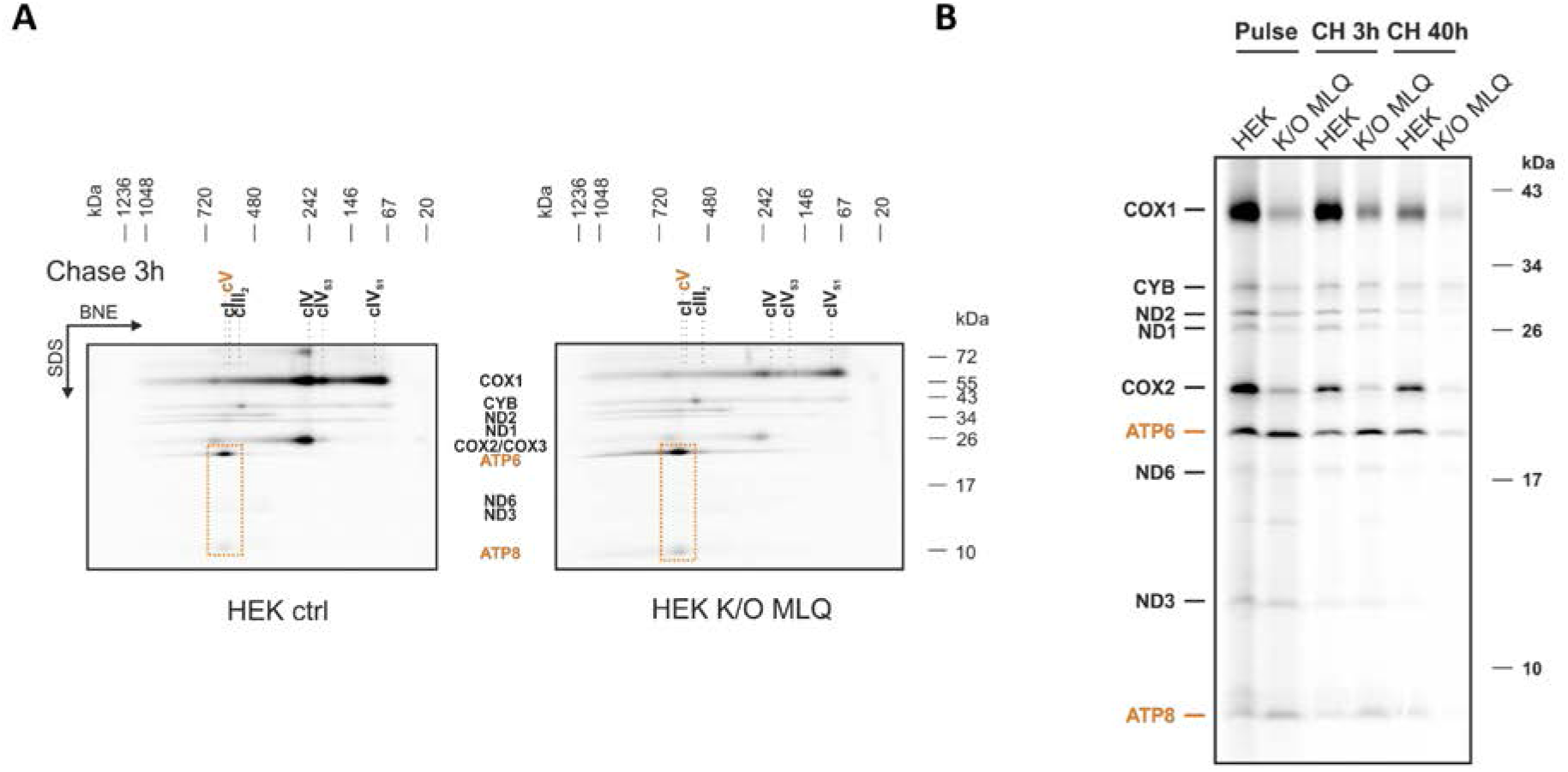
Supplementary to Fig. 5. **(A)** mtDNA-encoded subunits were radioactively labelled with ^35^S-methionin/cysteine and subsequently chased for 3h with cold methionine. Cells were solubilised with digitonin and analysed using 2D BNE/SDS PAGE. **(B)** 1D SDS-PAGE analysis of the same samples as in 2D gels in figure 5 and S1 used for signal quantification presented in Fig 5 B.

**Figure S2:**
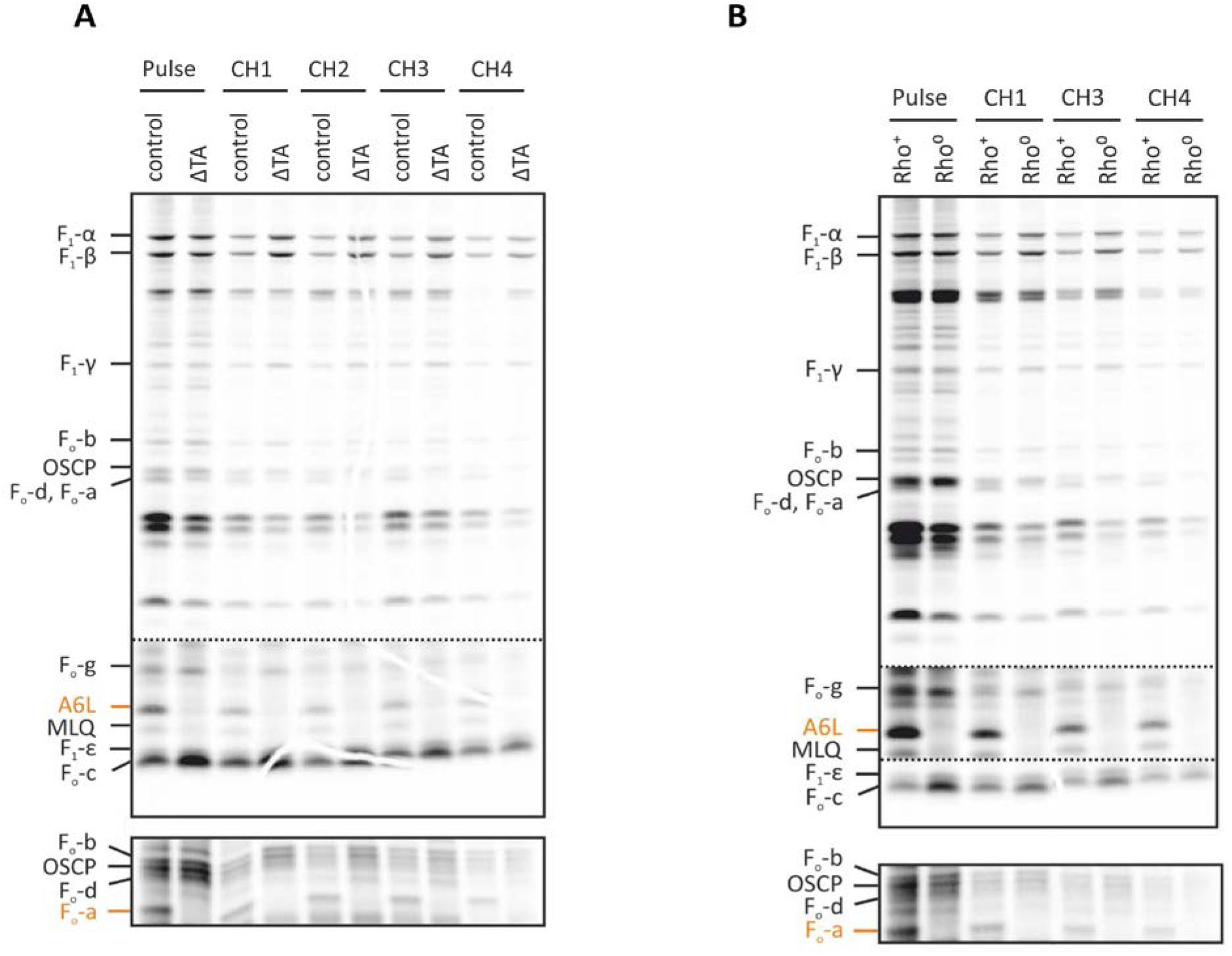
supplementary to Fig. 6. **(A)** ΔTA and control or (B) Rho^+^ and Rho^0^ cells were radioactively labelled for 3h with ^35^S-methionin/cysteine and subsequently chased with cold methionine for 1, 2, 3, and 4 days. Cells were harvested, solubilised with triton X-100 and ATP synthase was isolated by immune-affinity purification using ATP synthase immunocapture kit. IP-ATP synthase was than analysed by SDS-PAGE.

